# Mixed Heavy Metals Stress Induces Global Iron Starvation as Revealed by System Level Multi-Omic Analyses

**DOI:** 10.1101/2022.08.05.502853

**Authors:** Jennifer L. Goff, Yan Chen, Michael P. Thorgersen, Linh T. Hoang, Farris L. Poole, Elizabeth G. Szink, Gary Siuzdak, Christopher J. Petzold, Michael W.W. Adams

## Abstract

Globally, multiple heavy metal contamination is an increasingly common problem. As heavy metals have the potential to disrupt microbially-mediated biogeochemical cycling, it is critical to understand their impact on microbial physiology. However, systems-level studies on the effects of a combination of heavy metals on bacteria are lacking. Here, we use a native *Bacillus cereus* isolate from the subsurface of the Oak Ridge Reservation (ORR; Oak Ridge, TN, USA) — representing a highly abundant species at the site— to assess the combined impact of eight metal contaminants. Using this metal mixture and individual metals, all at concentrations based on the ORR site geochemistry, we performed growth experiments and proteomic analyses of the *B. cereus* strain, in combination with targeted MS-based metabolomics and gene expression profiling. The combination of eight metals impacts cell physiology in a manner that could not have been predicted from summing phenotypic responses to the individual metals. Specifically, exposure to the metal mixture elicited global iron starvation responses not observed in any of the individual metal treatments. As nitrate is also a significant contaminant at the ORR site and nitrate and nitrite reductases are iron-containing enzymes, we also examined the effects of the metal mixture on reduction of nitrogen oxides. We found that the metal mixture inhibits the activity of these enzymes through a combination of direct enzymatic damage and post-transcriptional and post-translational regulation. Altogether, these data suggest that metal mixture studies are critical for understanding how multiple rather than individual metals influence microbial processes in the environment.

## INTRODUCTION

Beginning in the 20^th^ century, increasing levels of heavy metal contamination occurred in both aquatic [1, 2] and terrestrial [3, 4] environments due to greater anthropogenic inputs from industrial and agricultural activities. Regardless of specific input source, a common theme that emerges at these contaminated sites is the simultaneous presence of multiple heavy metals at concentrations exceeding the threshold limits set by global environmental regulatory bodies [2, 5-8]. A recent meta-analysis by Zhou et al. (2020) compiled heavy metal concentrations for global surface water bodies and compared these values to the limits set by WHO and the US EPA. From 1972 to 2017, heavy metal pollution in surface waters shifted from single metal to multiple metal pollution.

Heavy metal pollution is not only detrimental to human [9], animal [10, 11], and plant health [12], it also has the potential to disrupt the natural cycling of elements via impacts on microbial activity. For example, a meta-analysis performed by Aponte et al. (2020) found that individual heavy metal contaminants linearly decreased the activities of key soil microbial enzymes, particularly those involved in carbon and sulfur cycling. In natural soil systems, individual heavy metal contaminants inhibit multiple steps of the denitrification pathways, resulting in accumulation of toxic intermediates, including nitrite and the greenhouse gas nitrous oxide [14-16]. However, studies investigating the impacts of a combination of, rather than a single, heavy metal on environmental microorganisms are scarce and have largely focused on determinations of IC_50_ values for binary combinations of metals rather than exploring systems-level impacts [17-23]. Nonetheless, these studies are informative as they demonstrate the potential for interaction between more than one metal. For example, Fulladosa et al. (2005) found that, in *Vibrio fisheri*, several heavy metal pairings were synergistic in their interactions, implying that the toxicity of these mixtures was greater than the sum of the toxicities of the individual metals [17].

The subsurface of the US Department of Energy (DOE) Oak Ridge Reservation (ORR) in Oak Ridge, Tennessee is contaminated with nitric acid and multiple heavy metals, making it an ideal for investigating the impacts of multi-metal contamination on native microbial communities [24]. The contamination is the result of liquid waste discharge from uranium processing operations at the Y-12 National Security Complex into on-site unlined waste ponds (referred to as the S-3 ponds) from 1951 until 1983 [25]. The subsurface of the region immediately adjacent to the former S-3 ponds, referred to as Area 3, is contaminated with high levels of uranium (U) and nitrate, as well as elevated concentrations of other metals, such as nickel (Ni), cadmium (Cd), copper (Cu), aluminum (Al), manganese (Mn), and iron (Fe) [25, 26]. As nitrate is a major co-contaminant, the impact of these metal contaminants on nitrogen cycling by microorganisms at the site is a significant point of concern [27, 28]. Previously, we isolated *Bacillus cereus* strain CPT56D-587-MTF (referred to as strain CPTF) from the subsurface sediments of the ORR Area 3. Strain CPTF carries out dissimilatory nitrate reduction to ammonium via NarGHI and NasDE [29]. Importantly, strain CPTF is a representative isolate of a highly abundant amplicon sequence variant (ASV) found in the Area 3 subsurface sediments. In a recent sediment survey, this ASV had the highest relative abundance across all Area 3 sediment samples with an abundance up to 10-40% of total reads in several samples, suggesting that strain CPTF represents a dominant taxon at the site [29]. Thus, this isolate is ideal for detailed studies of the physiological responses of native microbiota to site-relevant stressors.

We sought to assess the impact of a mixture of eight major metal contaminants of the Area 3 subsurface (Al, U, Mn, Fe, Cd, Cu, Co, and Ni) on the field isolate *B. cereus* strain CPTF. We compared the growth of strain CPTF in the presence of individual metals as well as in combination. Using a high-throughput MS-based proteomics approach, we compared the responses of cells exposed to the metal mixture to those exposed to individual metals. We further validated and expanded upon these results with a combination of targeted MS-based metabolomics, targeted gene expression profiling and enzyme activity assays. Finally, to explore the potential impact on nitrogen cycling at ORR Area 3, we examined the effects of the metal mixture on nitrate and nitrite reduction by strain CPTF.

## MATERIALS AND METHODS

### Media and culture conditions

*Bacillus cereus* str. CPT56D-587-MTF (referred to as strain CPTF) [30] was routinely streaked out on LB plates and grown overnight at 30 ºC. A single isolated colony was inoculated into LB broth and grown overnight shaking at 200 rpm at 30 ºC. The overnight culture was diluted 100-fold into an anoxic defined medium supplemented with nitrate (named *B. cereus* experimental medium or BCE medium) for further experimentation.

Detailed medium preparation procedures are in the **Supporting Information**. For metals exposure, a contaminated ORR environmental metal mix (COMM) [31] was used that contains:

500 µM AlK(SO_4_)_2_·12H_2_O, 50 µM UO_2_(CH_3_COO)_2_, 50 µM MnCl_2_·4H_2_O, 50 µM NiSO_4_·6H_2_O, 15 µM CoCl_2_·6 H_2_O, 5 µM (NH_4_)_2_Fe(SO_4_)_2_·6H_2_O, 5 µM CuCl_2_·2H_2_O, and 2.5 µM Cd(CH_3_COO)_2_·2H_2_O. The same metal salts and concentrations were used for the individual treatments.

### Growth curves

Strain CPTF was grown in 10 mL BCE medium amended with COMM or the individual metal concentrations described above. Growth was determined by taking optical density measurements at 600 nm (OD600) on a spectrophotometer.

### Proteomics analysis

Strain CPTF was grown in triplicate 50 mL cultures in anoxic BCE medium amended with COMM or the individual metal concentrations described above. Control cultures had no amendments. Following 10 hours of growth at 30 °C, samples were collected for proteomic analyses. Protein was extracted and tryptic peptides were prepared by following established proteomic sample preparation procedures detailed in the **Supporting Information**. Peptides were analyzed using an Agilent 1290 UHPLC system (Santa Clara, CA) coupled to a Thermo Scientific Obitrap Exploris 480 mass spectrometer (Waltham, MA). The LC-MS acquisition method and DIA-NN configurations for peptide identification and quantification are described in the supporting information. Quantitative matrices on the protein level were extracted from the main DIA-NN reports and processed by an automated python script described in the established protocol [32]. Differentially expressed proteins with statistical significance were reported.

### Metabolomic analysis

Quantitative mass spectrometry (MS)-based metabolomics was used to validate proteomic observations of differentially regulated metabolic pathways. Strain CPTF was grown in 200 mL BCE medium (n=5 replicates) amended with COMM. Control cultures had no further amendments to the medium. Following 10 hours of growth at 30 °C, the cells were sampled for analysis. Sample preparation and analytical conditions are described in the **Supporting Information**. Intracellular metabolites were quantified using an Agilent 6495 triple quadrupole mass spectrometer with a jet stream source, coupled to an Agilent 1290 UPLC stack. MS transition states for the targeted compounds are given in **Table S1**. Data were processed using Agilent Quantitative software. Limits of quantification are given in **Table S2**.

### Iron uptake

Strain CPTF was grown in triplicate in 500 mL BCE medium. Control cultures had no further amendments to the medium. A second set of control cultures was amended with 5 µM (NH_4_)_2_Fe(SO_4_)_2_·6H_2_O—the same amount of iron (Fe) present in the COMM. COMM-treated cultures were amended with metals as described above. Prior to inoculation, a medium sample was collected from each culture bottle for determination of initial Fe concentrations. Cultures were then grown for 10 hours at 30 °C and sampled for analysis. Inductively coupled plasma/mass spectroscopy (ICP-MS) analysis was performed using an Agilent 7900 single quadruple mass spectrometer to quantify the total iron content of the uninoculated culture medium and whole cell extracts. Sample preparation procedures and analytical methods are in the **Supporting Information**.

### Enzyme activity assays

For both nitrate and nitrite reductase activity assays, strain CPTF was grown in 500 mL of BCE medium with or without COMM addition at 30°C. After 10 hours of growth, cells were harvested by centrifugation at 6,600 x *g* at 4°C for 15 minutes. Cells were washed once in pre-chilled 50 mM potassium phosphate buffer (pH 7.0). Nitrate and nitrite reductase assays were performed using a modified version of the procedure described by Thorgersen and Adams (2016). Detailed protocols are in the **Supporting Information**.

### Quantitative reverse transcriptase PCR (qRT-PCR)

Strain CPTF was grown in triplicate 50 mL cultures in BCE medium. One set of cultures was left untreated as a control. One set was treated with the COMM described above. Following 10 hours of growth at 30 °C, RNA was extracted, and cDNA was prepared as described in the **Supporting Information**. Quantitative qRT-PCR was performed with the Brilliant II SYBR Green QPCR Master Mix (Agilent). Primers were designed to amplify ∼150 bp product within the target genes. The *recA* gene (UIJ64731.1) was used as the reference gene. Primer sequences are given in **Table S3**. Statistical comparisons were performed using a Student’s t-test.

## RESULTS AND DISCUSSION

### Growth of strain CPTF with a synthetic mixture of metals mimicking a contaminated site

In the ORR Area 3 subsurface, multiple metals co-exist at elevated concentrations along with high levels of nitrate [25]. To explore the toxicity of these metals, we exposed the ORR isolate *B. cereus* strain CPTF to a contaminated ORR environmental metal mix (COMM) containing eight metals, Al (500 µM), U (50 µM), Mn (50 µM), Fe (5 µM), Cd (2.5 µM), Co (15 µM), Cu (5 µM) and Ni (50 µM), where the concentrations reflect those typically found in ORR Area 3 groundwater **(Table S4)**. Compared to the control, strain CPTF cultures grown with COMM had a slower growth rate and lower growth yield **(Fig. 1A)**. To determine if the individual metals also inhibit growth, the cells were grown with the individual COMM components at the same concentrations they are present at in the COMM. For all the above-mentioned metal exposed cultures, we compared growth at the transition point between exponential and stationary phase (10 h) **(Fig. 1B)**. There was no significant difference (p > 0.05) between control culture growth and growth during Al, U, Mn, Co, Fe, or Cd. However, growth with Ni and Cu was 87 ± 3.9% and 92 ± 4.4% of the control, respectively (p < 0.05). We considered that the toxicity of the COMM may be the result of the additive effects of the two metals that were mildly toxic to the cells: Ni and Cu. If this were true, we would predict that growth of the COMM cultures would be 79 ± 5.9% of the control. However, the actual growth defect with COMM was more pronounced at 63 ± 4.8% of the control. These data suggest that the combined toxicity of the COMM metals is greater than what would be predicted from the sum of the individual parts.

**Figure 1.**
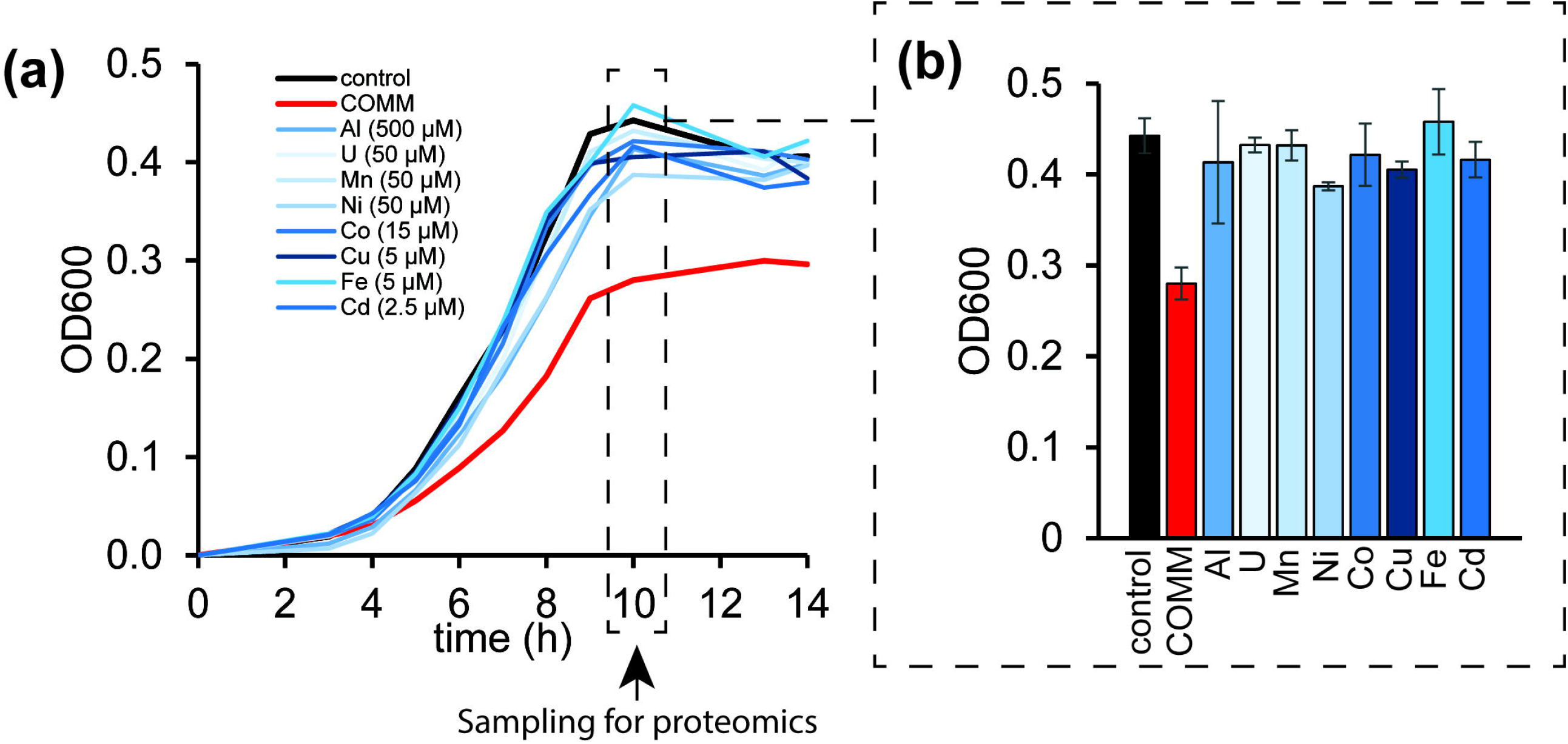
Growth of strain CPTF with field-relevant concentrations of individual metals or a metals mixture (COMM). **(a)** Growth assessment was performed under anoxic nitrate-respiring conditions with glucose (20 mM) as the carbon source. The added concentrations of the individual metals are indicated in the legend. The COMM is a mixture of all the metals at the same concentrations as they are present individually (**Table S4)**. Each time point represents an average of three replicates. For clarity, error bars are not shown but can be viewed in **Table S5**. The dashed box indicates the 10 h time point where samples were collected for the proteomic analysis described later in the manuscript. **(b)** Growth at the 10 h time point. Error bars represent ± SD. Data are the average of three replicates.

### MS-based proteomics analysis of metals-treated CPTF cultures: global response

We further examined the interactions of the COMM metals within the cellular system of strain CPTF using a proteomics-based approach. We compared the proteomic response of cells grown with COMM to cells grown with the individual metal treatments. Triplicate CPTF cultures were grown with either the COMM or individual metals at the same concentrations that they are present at in the COMM. Control cultures had no added metals other than those in the standard medium. Samples were collected for MS-based proteomics analysis at 10 h **(Fig. 1)**. Across all ten conditions a total of 1,303 unique proteins were identified **(Table S6)**. Differentially expressed proteins in metal-treated cultures were identified through comparison to control cultures without metal exposure. In sum, 295 significantly differentially expressed proteins (p < 0.05) were identified across the nine metal treatments (COMM and eight individual metals; **Fig. 2A, Table S7)**. Interestingly, of these 295 proteins, 191 (65%, termed Group 1) were uniquely differentially expressed in the COMM-treated cultures. The remainder (35%, Group 2) were differentially expressed in the COMM and one or more individual metal treatments or only in the individual metal treatments.

**Figure 2.**
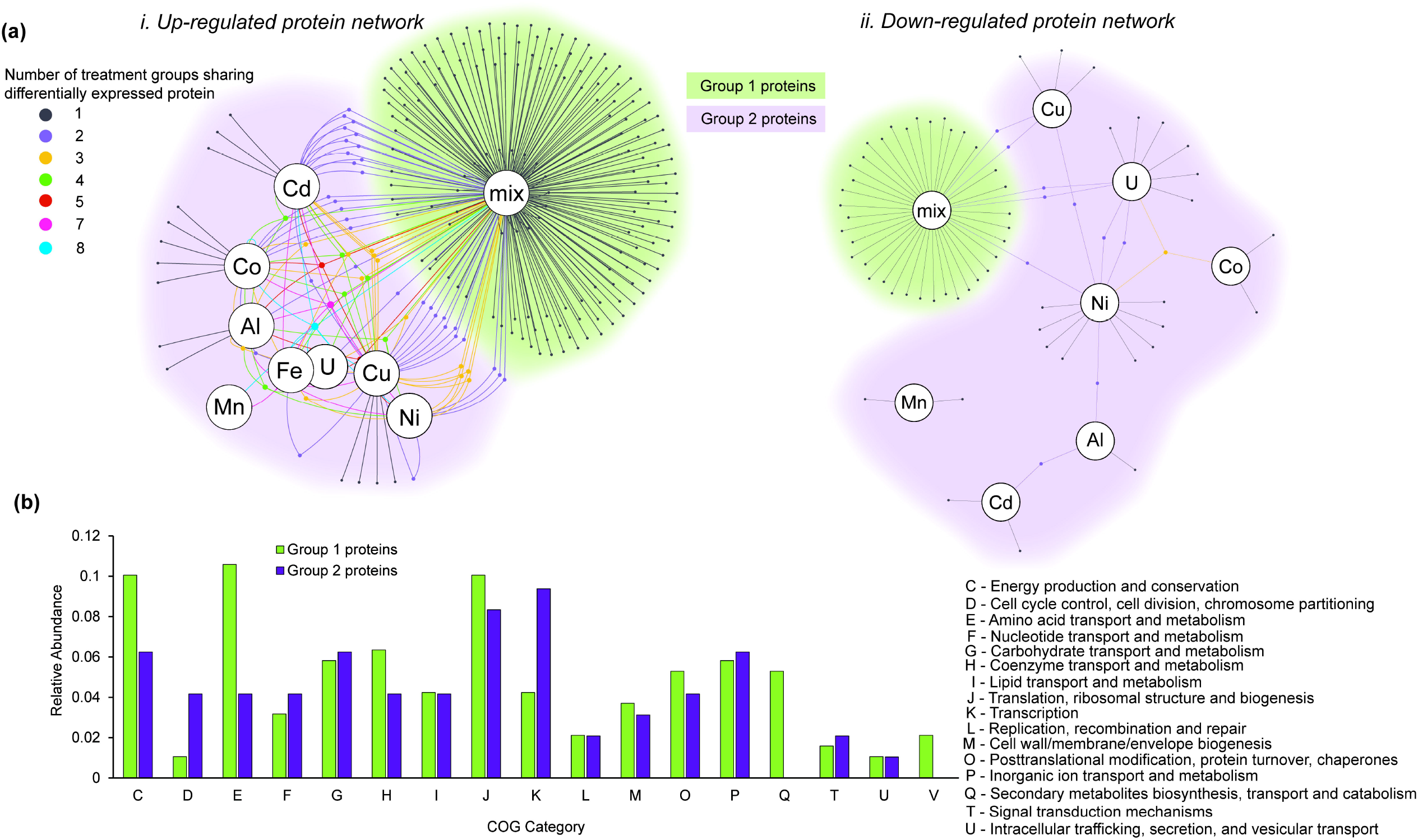
Comparison of the global response of the strain CPTF proteome between metals mix and individual metal treatments. **(a)** Network diagrams of differentially expressed proteins at 10 h of growth under anoxic nitrate-respiring conditions (n=3 replicates). Large nodes represent the metal treatment condition. Small nodes represent individual differentially expressed proteins. Edges indicate conditions under which the proteins are differentially expressed. Both significantly up-regulated (i) and down-regulated (ii) protein sets are displayed. Small node/edge colors (legend is shown on the image) represent the number of treatment groups sharing the differentially expressed proteins. Background cloud colors distinguish the Group 1 proteins (green: only expressed in the metals mix treatment) from the Group 2 proteins (purple: all other proteins). **(b)** Functional comparisons of Group 1 and Group 2 proteins using COG categories. Category S (unknown function) is excluded from the visualization. All other categories were not represented in the differentially-expressed proteomes.

### Competing siderophore and tryptophan biosynthesis pathways are differentially expressed only during COMM exposure

A total of 276 out of the 295 differentially expressed proteins were assigned to Clusters of Orthologous Genes (COG) categories [34] **(Fig. 2B)**. Functional comparison of Group 1 to Group 2 proteins revealed a ∼2.5-fold enrichment of COG Category E (amino acid metabolism and transport) among the Group 1 proteins: 11% of Group 1 proteins belonged to this category compared to 4% of Group 2 proteins. A more detailed analysis revealed that a number of these Group 1 Category E proteins were involved in biosynthesis of the essential amino acid tryptophan. Additionally, none of the Group 2 proteins belonged to Category Q (secondary metabolites biosynthesis, transport, and catabolism). In contrast, ten (5%) of the Group 1 proteins fell into COG Category Q. These included several involved in the biosynthesis of the siderophore bacillibactin. Bacillibactin is a tripeptide catecholate-type siderophore secreted by members of the genus *Bacillus* for chelation of ferric iron (Fe^3+^) from the environment [35].

In concert, these two observations are interesting as both tryptophan and bacillibactin biosynthetic pathways of strain CPTF are predicted to share chorismite as intermediate [36, 37]. Chorismate is the branch point in the biosynthesis of aromatic amino acids and other aromatic metabolites, such as siderophores [38]. Further analysis of specific protein expression patterns revealed that the bacillibactin biosynthesis enzymes isochorismate synthase (DhbC), 2,3-dihydrozybenzoate dehydrogenase (DhbB), and (2,3-dihydrozybenzoate)adenylate synthase (DhbE) are significantly up-regulated in response to COMM treatment but not to exposure to individual metals **(Fig. 3A)**. Where commercial standards were available, we performed targeted MS-based analysis of intracellular metabolites to confirm changes in metabolic pathway flux suggested by the protein expression data. DhbC (+6.8 log_2_FC) catalyzes the conversion of chorismate to isochorismate. Likely due to its role as a branch point intermediate, chorismate concentrations were low in all samples and no difference was observed in its concentrations between control and COMM cultures **(Fig. 3B)**. Isochorismate is then converted to 2,3-dihydroxy-2,3-dihydroxybenzoate by DhbB (+9.6 log_2_FC), and then to 2,3-dihydroxybenzoate by an enzyme that was not differentially expressed. We found that intracellular 2,3-dihydroxybenzoate levels increased 8.7-fold in response to COMM exposure **(Fig. 3B)**, likely due to greater metabolic flux through the pathway. In bacteria such as *Bacillus subtilis*, 2,3-dihydroxybenzoate is secreted as a siderophore in addition to its role as a biosynthetic intermediate [39, 40]. Adenylation of 2,3-dihydrozybenzoate to (2,3-dihydrozybenzoate)adenylate is catalyzed by DhbE (+6.4 log_2_FC). The final step is the formation of the tripeptide bacillibactin mediated by a non-ribosomal peptide synthase not differentially expressed in our dataset.

**Figure 3.**
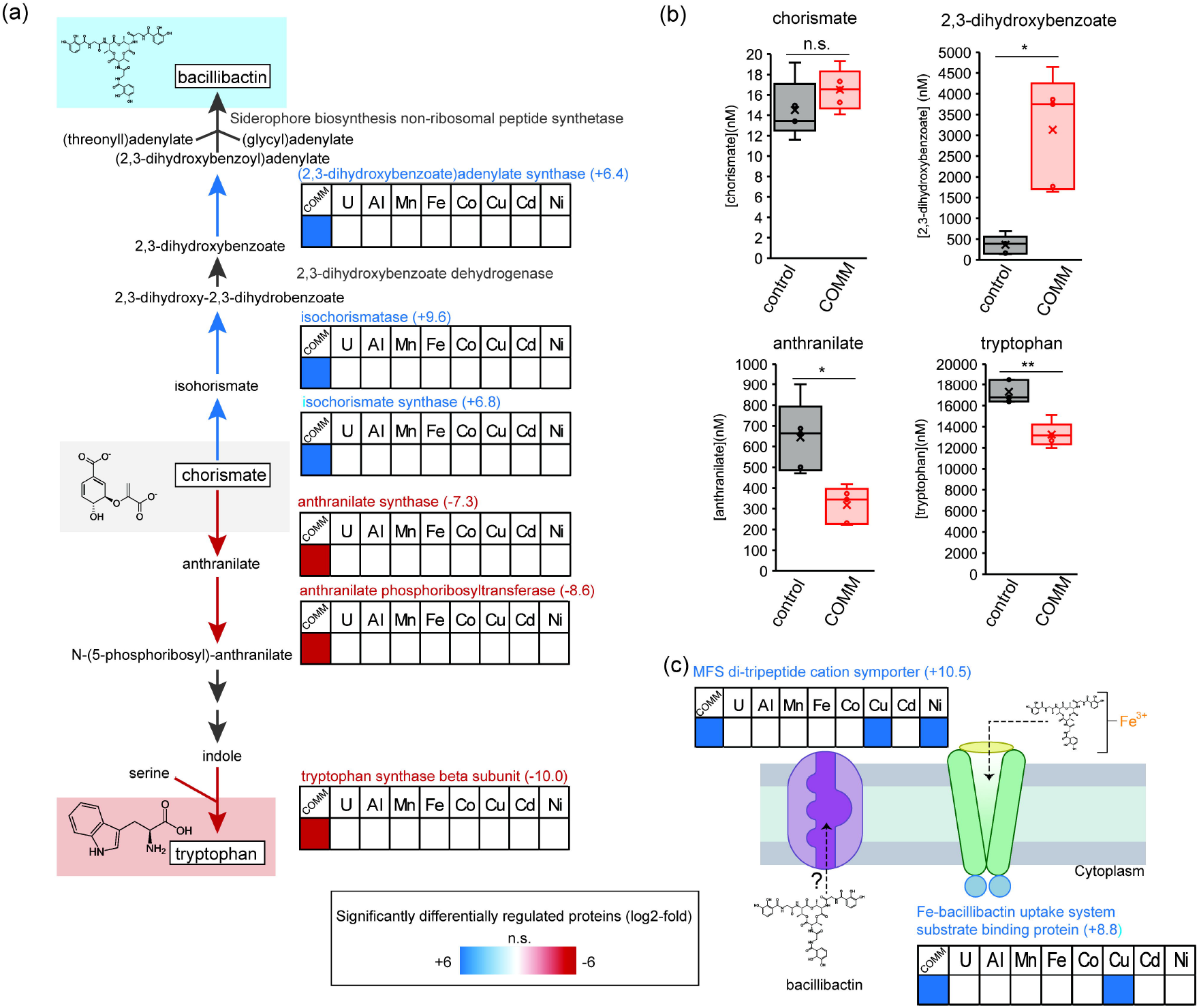
COMM exposure dysregulates bacillibactin and tryptophan biosynthetic pathways. **(a)** Protein expression patterns across the enzymes of the bacillibactin and tryptophan biosynthetic pathway. Heat maps display average (n=3 replicates) log_2_-fold expression changes of individual proteins across the different treatment conditions relative to the control (left-to-right: COMM, U, Al, Mn, Fe, Co, Cu, Cd, Ni). Numbers in parentheses to the right of protein names indicate the specific log_2_-fold change of the protein in the COMM-exposed cultures. The heat map scale (displayed at log_2_-fold expression changes relative to the control) is at the bottom of the image. Blue boxes indicate significantly up-regulated proteins (p<0.05), red boxes indicate significantly down-regulated proteins (p<0.05) and white boxes indicate no significant difference in expression relative to the control. Pathway proteins lacking a heat map had no significant change in expression patterns in any of the tested conditions. **(b)** Plots of intracellular concentrations of select bacillibactin/tryptophan biosynthetic pathway metabolites (n=5 replicates). Center lines represent median values. Interquartile ranges and maximum/minimum values are represented by box ranges and whisker ranges, respectively. Mean values are indicated with an “x”. Internal points are marked with dots. * p < 0.01, ** p < 0.001. **(c)** Protein expression patterns of putative bacillibactin transporters. Heat map details are the same as panel **(a)**.

In contrast to the results for bacillibactin biosynthetic pathway of strain CPTF, expression of the three enzymes in the tryptophan pathway after chorismate were significantly down-regulated in response to COMM treatment **(Fig. 3A)**, although there was also no change in expression with any of the individual metal treatments. These down-regulated proteins include anthranilate synthase (TrpE, -7.3 log_2_FC), which catalyzes chorismate conversion to anthranilate. Supporting this observation, intracellular anthranilate concentrations decreased 2.0-fold in response to COMM exposure **(Fig. 3B)**. Anthranilate phosphoribosyltransferase (TrpD), which catalyzes conversion of anthranilate to N-(5-phosphoribosyl)-anthranilate, was also downregulated (−8.6 log_2_FC). Finally, down-regulation of tryptophan synthase beta subunit (TrpB, -10.0 log_2_FC) was observed. TrpB catalyzes the conversion of the downstream intermediate indole to tryptophan. As a result, intracellular tryptophan concentrations decreased 1.3-fold during COMM exposure **(Fig. 3B)**.

We also observed increased expression of two putative siderophore transporters during COMM exposure, although we note that these transporters were part of the Group 2 proteins as they were also expressed during some individual metal treatments **(Fig. 3C)**. The substrate-binding subunit (FeuA, +8.8 log_2_FC) of the FeuABC bacillibactin-Fe^3+^ ATP-type transporter [41] was significantly up-regulated during COMM exposure. In addition, while there is no homolog of a characterized bacillibactin exporter [42] in the strain CPTF genome, we observed increased expression (+10.5 log_2_FC) of a major facilitator superfamily (MFS) type di/tripeptide cation symporter during growth with COMM (as well as individual Cu and Ni exposure). The previously characterized bacillibactin transporter of *B. subtilis* (YmfE) is also of the MFS-type [42]. Further supporting this proposed function, the gene encoding this transporter (LW858_01095) is located immediately adjacent to a second ABC-type siderophore-Fe^3+^ transport system (**Fig. S1)** and is downstream of a metal-responsive ArsR-type transcriptional regulator [43].

Our proteomic and metabolomic data suggest an increase in bacillibactin and/or 2,3-dihydrozybenzoate production, export and re-import occurs during COMM exposure of *B. cereus* str. CPTF. However, the biosynthesis and membrane transport of siderophores are energetically costly processes [44, 45]. Additionally, iron uptake is typically tightly-regulated by bacterial cells to prevent mismetallation of metalloproteins [46] and oxidative damage triggered by excessive intracellular iron [47]. In *B. subtilis* str. CU1065 and *B. cereus* str. 569, the genes encoding siderophore biosynthetic enzymes and transporters are regulated by a canonical ferric uptake regulator (FUR) that represses its regulon under iron-replete conditions [48, 49]. During periods of iron starvation, the FUR regulon is derepressed. We propose that COMM exposure induced a global iron starvation response in the cell involving increased siderophore biosynthesis as well as increased expression of siderophore transporters. Additionally, we suggest that chorismate is shunted from tryptophan to bacillibactin biosynthesis for prioritization of iron acquisition by strain CPTF.

### COMM exposure results in a canonical global iron starvation response

We examined the protein expression data and conducted targeted gene expression analyses to determine if the alterations observed in the bacillibactin and tryptophan pathways were indicative of a more global iron starvation response to COMM exposure **(Table 1)**. For these analyses, we compared our expression data to prior studies of cellular responses to iron limitation conditions. We confirmed the presence of a FUR homolog in the strain CPTF genome with 83% sequence identity (full length) to *B. subtilis* FUR. In *B. subtilis*, as well as other model microorganisms, this FUR-regulated response includes: (1) increased expression of transporters for iron uptake (e.g. siderophore-Fe^3+^ transporters) and iron scavenging proteins, (2) siderophore biosynthesis, and (3) an iron-sparing response including increased expression of flavodoxins [48, 50]. We have presented above evidence for increased iron transport in strain CPTF. Related to iron-scavenging, we note the increased expression of two heme monooxygenases (HmoA and HmoB) exclusively during COMM exposure **(Table 1)**. HmoA and B catalyze heme porphyrin ring opening to release free Fe^2+^ [51]. Heme monooxygenases are expressed by *Bacillus* and *Staphylococcus* species during periods of iron limitation [52, 53].

**Table 1.**
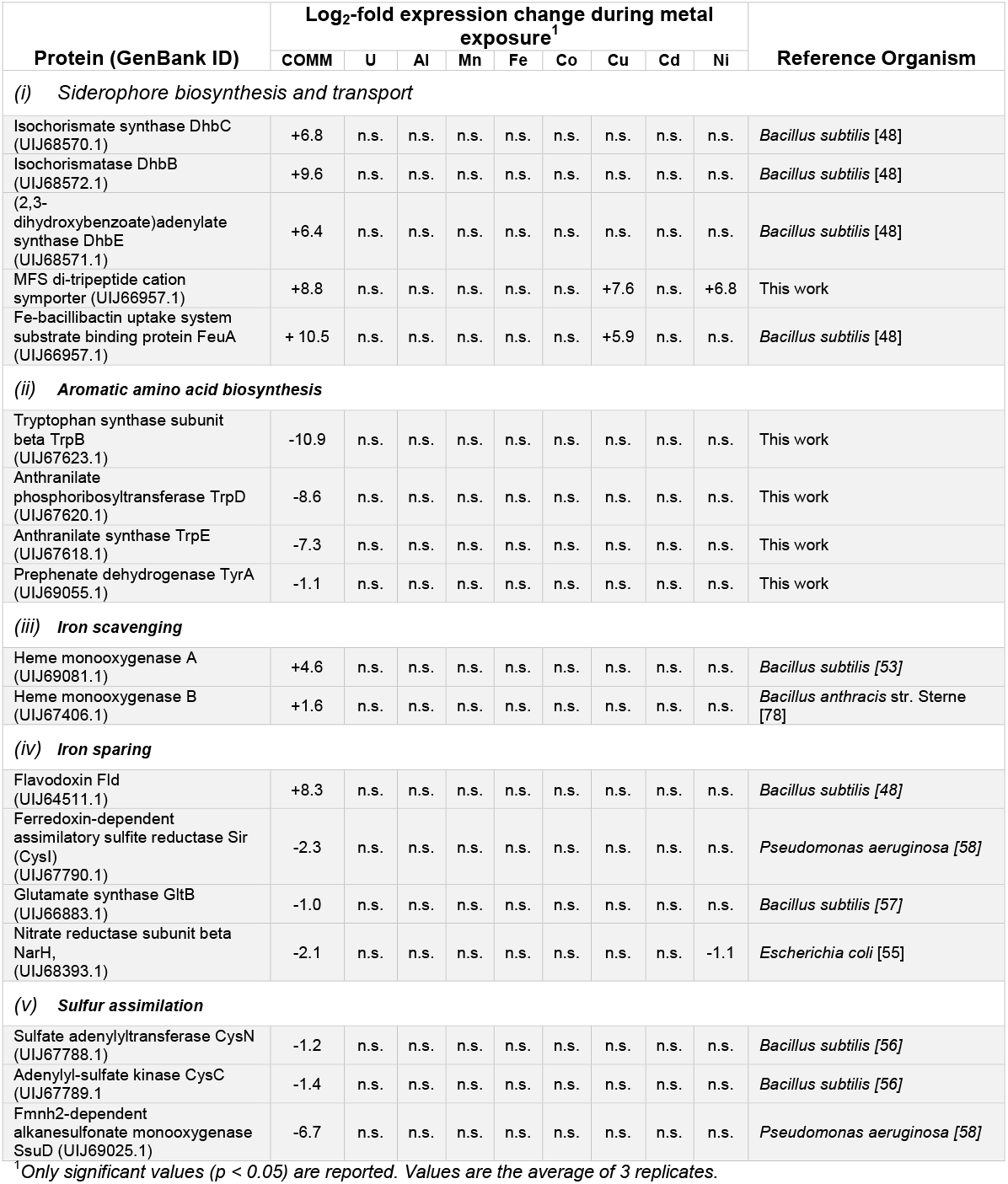
Changes in expression levels of proteins proposed to comprise the strain CPTF global iron starvation response.

As discussed above, we observed increased expression of genes for bacillibactin biosynthesis, consistent with what has been observed with other *Bacillus* species during periods of iron starvation [48, 49]. While we suggest that the expression changes in the tryptophan pathway are for conservation of chorismite, a key biosynthetic intermediate, we were unable to find any reported instance of tryptophan biosynthesis repression in the global iron starvation response of other bacterial species. We propose that this response may represent novel regulatory adaptation of strain CPTF to prioritize metabolic intermediates for iron acquisition during the extended periods of heavy metal stress in its natural environment at the ORR. In further support of this model, we observed decreased expression of the prephenate dehydrogenase (TyrA) exclusively during exposure to COMM and not to individual metals **(Table 1)**. TyrA catalyzes the oxidative decarboxylation of prephenate to 4-hydroxyphenylpyruvate in the tyrosine biosynthetic pathway. Like tryptophan biosynthesis, this pathway utilizes chorismate as an intermediate [54]. Future studies are required to determine how this response is regulated.

To respond to iron-limited conditions, bacteria typically have an iron-sparing response to reduce synthesis of abundant, non-essential, iron-bearing enzymes and enzymes involved in iron cofactor biosynthesis [55]. The iron-sparing response of *B. subtilis* is under post-transcriptional regulatory control of the sRNA FsrA (analogous to RhyB in gram-negative bacteria) and three small, basic proteins (FbpA, FbpB, and FbpC) [56]. This response includes decreased expression of iron-containing enzymes of the TCA cycle, cytochrome and porphyrin biogenesis enzymes, cysteine biogenesis enzymes, iron-containing enzymes of the isoleucine biosynthetic pathways, and iron-containing glutamate synthase [56, 57]. During COMM exposure of strain CPTF, we similarly observed decreased expression of the sulfate adenylyltransferase (CysN) and the adenylyl-sulfate kinase (CysC) as well as the ferredoxin-dependent sulfite reductase (Sir/CysI) **(Table 1)**. We speculate that the decreased expression of the non-iron containing enzymes of the cysteine biosynthetic pathway is due to the role of cysteine as the sulfur donor in iron-sulfur cluster biosynthesis [58]. We also observed decreased glutamate synthase (GltB) levels during COMM exposure **(Table 1)**. Finally, we note decreased expression of the NarH subunit of the respiratory iron-sulfur cluster-containing nitrate reductase. Prior studies with *B. subtilis* under iron-limited conditions were conducted under aerobic conditions when respiratory nitrate reductease is not produced but decreased expression of this enzyme has been shown in *Escherichia coli* during iron limitation under anaerobic conditions [59].

Connected to the iron-sparing response, the protein flavodoxin (Fld) is upregulated across all three domains of life during periods of iron limitation [60-62]. Flavodoxins are small electron transfer proteins that contain a flavin mononucleotide (FMN) cofactor. During low-iron conditions, organisms will up-regulate expression of flavodoxin and this replaces iron-sulfur containing ferredoxin as the physiological electron donor for various oxidoreductase reactions [60, 62]. Accordingly, we observed increased production of flavodoxin in the COMM-exposed proteome of strain CPTF relative to the control (**Table 1**). In contrast, flavodoxin was not up regulated by any of the individual metal treatments.

We used qRT-PCR to validate representative protein expression changes observed by proteomics during COMM exposure of strain CPTF. We selected 11 transcripts to represent the various parts of the proposed global iron starvation response of strain CPTF. Of these 11 transcripts, seven (*dhbB, dhbC, dhbE, mfs, sir, fld*, and *hmoA*) matched the expression patterns observed for their respective protein products during COMM exposure **(Fig. S2)**. However, four transcripts (*feuA, trpB, trpD*, and *trpE*) were not differentially expressed despite large changes in their protein product abundances observed under the same conditions. The discrepancy between the *feuA* transcript expression levels (n.s.) and FeuA protein expression levels (+8.8-log_2_FC) is likely the result of temporal changes in gene expression during the transition into stationary phase that begins at our sampling time-point (10 h) and subsequent lag in protein-level changes [63]. In contrast, TrpBDE protein expression is decreased during COMM exposure while its transcript abundance is unchanged, suggesting that post-transcriptional or post-translational regulation. These results emphasize the utility of combined proteomic and transcriptomic approaches.

While increased expression of proteins involved in siderophore biosynthesis and transport has been observed previously during individual metal exposure experiments [64, 65], there are no reports of an iron-starvation response occurring to the extent observed here with strain CPTF during COMM exposure. Additionally, these prior studies have either utilized high heavy metal concentrations that are not relevant even to contaminated environments, or the heavy metal exposure is performed under iron-deficient conditions [66]. For example, in *B. subtilis*, copper exposure (500 µM) induces expression of bacillibactin biosynthetic enzymes as well as the bacillibactin-Fe^3+^ transporter. However, no expression changes were observed for the heme monooxygenases or flavodoxin. Furthermore, several iron-containing enzymes were actually up-regulated during copper exposure and there was no evidence of an iron-sparing response [67]. Thus, exposure of strain CPTF to COMM appears to induce a physiological state of iron starvation comparable to that induced by iron chelators or what is observed in Δ*fur* mutant strains [48].

### Proposed mechanism for iron starvation response in COMM-exposed cells

Iron concentrations are elevated in the highly contaminated ORR groundwater ([Fe]_AVG_ = 5.6 µM) relative to non-contaminated groundwater ([Fe]_AVG_ = 0.35 µM) at the site [28, 31]. In our experiments, [Fe] was 3.7 and 5.3 µM Fe in the control and COMM-amended cultures, respectively. The [Fe] in the control is higher than in the non-contaminated groundwater due to the iron present in the base medium plus contaminating iron from other medium components. Importantly, [Fe] in the COMM cultures is nearly the same as the average for the contaminated groundwater. In the culture medium, iron was always added in the ferrous form. Nonetheless, COMM exposure induces a physiological state of iron starvation that is not observed in the individual metal treatments or the controls. We considered that COMM components may compete with Fe^2+^ for the same transporter, limiting Fe^2+^ uptake. However, compared to unamended strain CPTF cultures, we found that COMM-exposed cultures over-import iron with total intracellular iron concentrations of 22.0 and 151.1 µM, respectively **(Fig. S3)**. As an additional control, we amended CPTF cultures with 5 µM ferrous iron (the same as that present in the COMM) and measured intracellular iron concentrations. While these cultures do take up about 50% more iron than the unamended control (33.6 vs. 22.0 µM, respectively), these values are still less than that measured for the COMM-exposed cultures.

Individual metals within the COMM may also disrupt intracellular iron homeostasis by displacing iron in the metal-binding sites of enzymes, a process known as mismetallation, which is a known mechanism of toxicity for many heavy metals [68]. We propose that individual metals in the COMM may target different stages of the iron cofactor assembly and insertion processes for different enzymes. During individual metal exposure, the cells appear to manage this stress by up-regulating proteins for iron cofactor repair and biosynthesis with minimal or no growth defect (**Fig. 1)**. For example, we observed increased expression of iron-sulfur cluster repair and assembly proteins (IsA and Ric) as well as heme biosynthesis proteins (HemFH) during exposure of strain CPTF to Cd and Cu **(Fig. S3)**. However, in combination, that stress response is seemingly overwhelmed, perhaps as multiple metals target a broader range of iron-bearing enzymes, resulting in significant disruption of intracellular iron homeostasis. Interestingly, the levels of two heme biosynthesis proteins (HemFH) are increased during COMM exposure relative to the control while the iron-sulfur cluster repair/assembly proteins are not, suggesting that iron may be prioritized for heme biosynthesis under these conditions. Increased rates of iron cofactor synthesis required to overcome the displacement of iron by other metals could lead to a significant shift in the intracellular Fe^2+^ equilibrium, leading to FUR de-repression.

### Potential impacts of COMM-disrupted iron homeostasis on nitrogen cycling

Nitrate-respiring conditions dominate in the ORR Area 3 subsurface environment due to the high concentrations of nitrate present as a major component of the contamination plume.

However, nitrogen cycling at the site may be influenced by heavy metal co-contaminants. Notably, nitrite accumulation has been measured in the porewaters of contaminated ORR sediment cores, suggestive of inhibition of nitrite reduction *in situ* [69]. The NarGHI respiratory nitrate reductase and the NasDE nitrite reductase both contain iron cofactors [70, 71]. Thus, we sought to determine if the dysregulation of iron homeostasis in strain CPTF induced by COMM impacts the activity of these two enzymes. We found that the activity of the NasDE nitrite reductase was near-absent (0.25 ± 0.44 units·OD600^-1^) in COMM-exposed CPTF cultures relative to the control (12.11 ± 7.48 units·OD600^-1^) **(Fig. 4A)**. However, expression levels of both protein subunits were unchanged between control and COMM-exposed cultures, **(Fig. 4B)**, suggesting that the loss of activity is due to direct inhibition of the enzyme by the metals.

**Figure 4.**
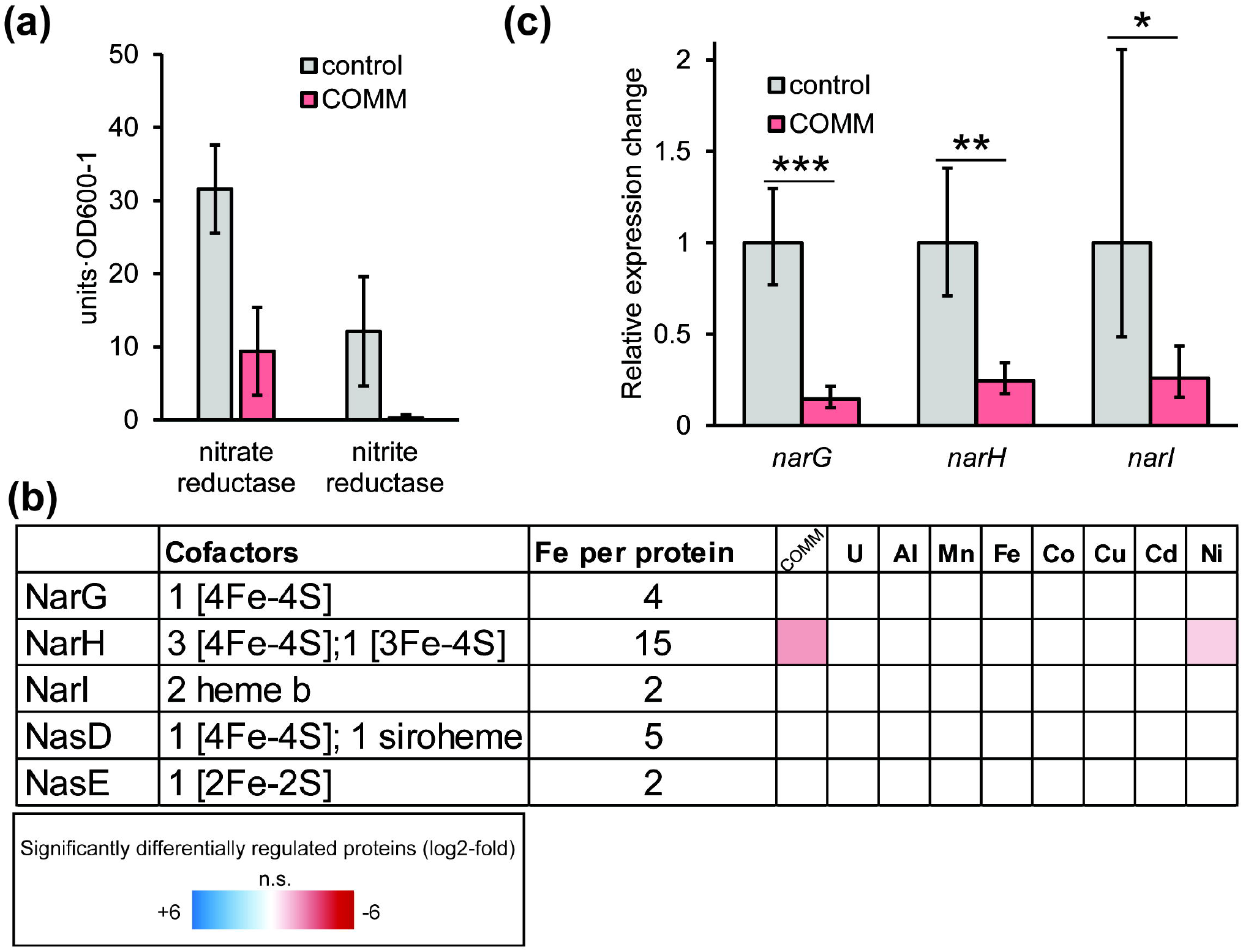
COMM treatment impacts nitrogen oxide reduction activity. **(a)** Activity of nitrate and nitrite reductases with (red bars) or without (grey bars) COMM. One unit of activity represents 1 nmol nitrate/nitrate reduced per minute. Experiments were performed in triplicate and error bars represent SD. **(b)** NarGHI and NasDE expression patterns. Heat map display average (n= 3 replicates) log_2_-fold expression changes of individual proteins across the different treatment conditions relative to the control (left-to-right: COMM, U, Al, Mn, Fe, Co, Cu, Cd and Ni). The heat map scale (displayed at log_2_-fold expression changes relative to the control) is at the bottom of the image. The number of Fe atoms per protein was determined from their *Bacillus subtilis* (UP000001570) homologs **(c)** Relative changes in expression of *narG, narH, and narI* with (red bars) and without (grey bars) COMM exposure. Experiments were performed in triplicate and error bars represent ±SD. * p < 0.05, ** p < 0.01, *** p < 0.001.

We also observed decreased nitrate reductase activity in the COMM-exposed strain CPTF cultures relative to the control (9.38 ± 6.00 v. 31.57 ± 6.03 units·OD600^-1^) **(Fig. 4A)**. As noted above, COMM exposure resulted in decreased levels of the NarH subunit of the nitrate reductase compared to the control **(Fig. 4B)**, suggesting that the decrease in activity is, in part, due to lower protein levels. Interestingly, no changes were observed in NarG and NarI expression **(Fig. 4B)**. This difference in expression levels changes between the three subunits is puzzling as all three are present in the same operon and should be co-regulated. Indeed, gene expression analyses confirmed the decreased expression of *narH* as well as *narI* and *narG* (**Fig. 4C)**. This discrepancy between the protein-level and transcript-level fold-changes is likely due to different post-translational controls on the three protein products. The observed decreased *narGHI* expression is likely part of the iron-sparing response described above. At a later timepoint, all three protein subunits would likely have decreased in the COMM-exposed cultures relative to the control. However, at the time point where all measurements were conducted, NarH levels may have already been subject to post-translational regulation, possibly via enhanced proteolytic degradation. Mismetallation of metal cofactor centers can result in misfolding, targeting proteins for cleavage by proteases [72]. Interestingly, we observed increased expression of two CPTF proteases exclusively during COMM treatment: Clp and Hls. Notably, NarH contains a greater number of iron atoms than either NarG or NarI: NarH contains a predicted 15 iron atoms per protein molecule compared to 4 and 2, respectively **(Fig. 4B)**. We speculate that this may make NarH more vulnerable to mismetallation and subsequent turnover than the other subunits. An integrated model for the impacts of COMM on nitrate and nitrite reduction is presented in **Figure 5**. Significantly, our data suggest that the inhibition of nitrite/nitrate reductase activities frequently observed at heavy metal contaminated sites [15, 73], including ORR Area 3, may occur at multiple regulatory levels.

**Fig. 5.**
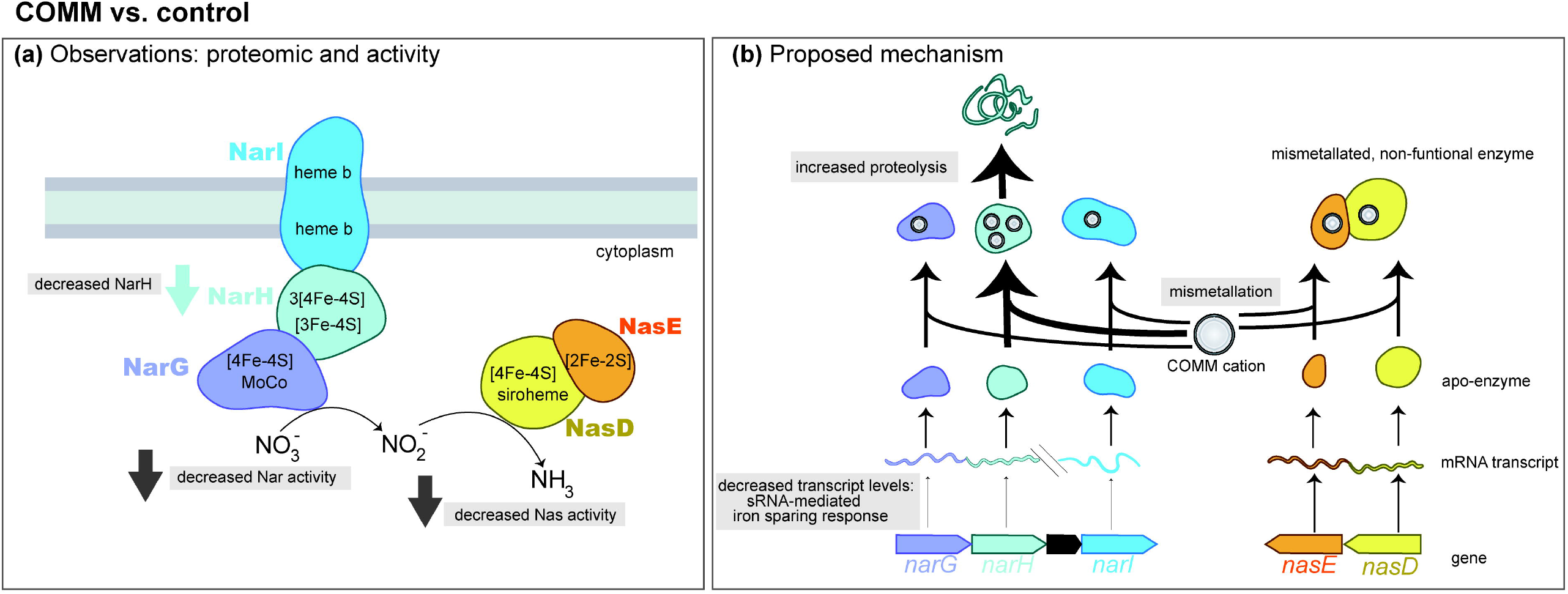
Model for decreased nitrate/nitrite reductase activity during COMM exposure. (a) Observed changes in enzyme levels and functions in COMM-exposed CPTF cells relative to the control. (b) Proposed mechanisms for changes described in panel (a).

### ECOLOGICAL IMPLICATIONS

Considering our findings, we searched previously published ORR metagenome data to determine if there is evidence of selection for iron acquisition-related genes within the heavy metal contaminated Area 3 subsurface that the COMM is designed to mimic. Interestingly, we found that Hemme et al. (2015) reported a significant enrichment of *efeU*, which encodes a high affinity ferrous iron transporter, in a groundwater metagenome from an Area 3 well (FW106, [U] = 160 µM, [Fe] = 0.8 µM, pH 3.6 [31]) relative to a groundwater metagenome from a pristine background well (FW301, [U] =0.004 µM, [Fe] = 0.07 µM, pH 7 [31]). EfeU is important for iron acquisition under iron-depleted conditions [75]. While other iron acquisition-related genes were not reported in this study, we would not expect to see enrichment for genes involved in siderophore production as many are highly conserved across diverse bacteria [76]. Likewise, the iron-sparing response observed here for ORR strain CPTF would not be apparent from metagenome data. Thus, meta-transcriptomic and meta-proteomic studies are called for to assess the differential expression of these systems at contaminated sites.

More importantly, our findings on metal-metal interactions in the cellular stress response of ORR strain CPTF, if validated in other microbial systems, have significant implications for how systems biology studies on metal toxicity are extrapolated to microbial processes occurring at contaminated environments. The construction of biological regulatory and metabolic networks for such sites may have limited predictive power if relying exclusively on data produced from single metal perturbation experiments as these models would underestimate the toxicity of the combined metals and extent of the systemic response.

## DATA AVAILABILITY

The generated mass spectrometry proteomics data have been deposited to the ProteomeXchange Consortium via the PRIDE [77] partner repository with the dataset identifier PXD035730, subject to a pre-publication embargo period. DIA-NN is freely available for download from https://github.com/vdemichev/DiaNN.

## ACKNOWLEDGEMENTS

We thank Lauren Lui and Torben Nielsen (Lawrence Berkeley National Lab) for initial assistance with searching the previously published metagenome data. JLG also thanks Nathan Yee (Rutgers) for the spirited discussions on the topic of metal resistance in the lab versus the field that inspired this work. This material by ENIGMA (Ecosystems and Networks Integrated with Genes and Molecular Assemblies) (http://enigma.lbl.gov), a Science Focus Area Program at Lawrence Berkeley National Laboratory, is based on work supported by the U.S. Department of Energy, Office of Science, Office of Biological and Environmental Research, under contract DE-AC02-05CH11231.

