## Supporting Information for "Mixed Heavy Metals Stress Induces Global Iron Starvation as Revealed by System Level Multi-Omic Analyses"

### CONTENTS:

**Supplementary Methods**……………………………………………………………………………….2
**Table S1.** MS transition states for metabolite analyses……………………………………..…….6
**Table S2.** Quantitation range for analyzed metabolites……………………………………..….7
**Table S3.** qRT-PCR primer sequence**s**…………………………………………………………..……8
**Table S4.** Geochemical parameters used to determine laboratory test values for experimentation with strain CPTF…………………………………………………………………………9
**Table S5.** Error bars (± SD) for Figure 1A…………………………………………………………10
**Table S6** CPTF full protein list…………………………………………………...See spreadsheet
**Table S7** CPTF all differentially expressed proteins heatmap…………………....See spreadsheet
**Figure S1.** Genomic location of the predicted bacillibactin exporter……………………….......11
**Figure S2.** qRT-PCR validation of protein expression changes……………………………...…12
**Figure S3.** Experimental insights into possible mechanism for iron starvation response……………………………………………………………………………………………...13
**References**…………………………………………………………………………………………14

### SUPPLEMENTARY METHODS

**Growth conditions.** This BCE growth medium contained, per liter: 2.5 g $NaHCO_3$, 0.6 g $NaH_2PO_4$, 0.5 g casamino acids, and 40 mL of a 25X stock solution of salts composed of 5 g/L $NH_4Cl$, 2.5 g/L KCl, 12.32 g/L $MgSO_4 \cdot 7H_2O$, 0.367 g/L $CaCl_2 \cdot 2\ H_2O$, and 0.25 g/L NaCl. Additionally, the medium contained, per liter, 1 mL of 1000X vitamin mix and 10 mL of a 100X DL minerals stock solution as described previously by Widdel and Bak [1]. The medium was routinely supplemented with 20 mM glucose and 10 mM $NaNO_3$ as a carbon source and terminal electron acceptor, respectively. The pH was adjusted to 7.2 using either HCl or NaOH. For anaerobic cultivation, the medium was purged of oxygen with 80%/20% $N_2/CO_2$. The cultures were incubated at 30 ºC.

**Quantitative metabolomics: sample preparation and analytical conditions.** CPTF cell pellets were collected by centrifugation at 4°C at 5500 x *g* for 15 minutes. The supernatant was discarded, and the pellet was resuspended in 5 mL of 1X PBS. The cell suspension was centrifuged again at 4°C at 5500 x *g* for 15 minutes. This wash step was repeated a second time. The cell pellet was then resuspended in 500 µL of 1X PBS and transferred to a microfuge tube and spun at 20,000 x *g* for 5 minutes. The supernatant was removed, and the pellet was spun down a second time to remove remaining liquid. The cell pellets were stored at -80 °C until extraction and analysis. For extraction of intracellular metabolites, 1000 µL of a cold methanol:acetonitrile:water solvent mixture (2:2:1, v/v/v) was added to the cell pellets followed by vortexing for 30 seconds. The samples were immediately submerged in liquid nitrogen for 1 minute. The samples were then thawed at room temperature and sonicated in an ice-cold water bath for 5 minutes. This freeze-thaw procedure was repeated two times. The samples were then incubated for 1 hour at -20 ºC followed by centrifugation at 16,000 x *g* at 4 ºC for 15 minutes. Afterwards, the supernatant was transferred to a clean tube and evaporated to dryness in a vacuum concentrator (LABCONCO CentriVap Benchtop). The dry extracts were reconstituted in 100 µL of a water:acetonitrile solvent mixture (1:1, v/v) and sonicated for 5 minutes in an ice-cold water bath followed by centrifugation for 15 minutes at 16,000 x *g* at 4 ºC to remove any insoluble debris. The supernatants were transferred to HPLC vials with inserts and stored at -80 ºC until LC/MS analysis.

For UPLC-based separation of the intracellular metabolites, the mobile phase was composed of A = 0.1% formic acid in water, and B = 0.1% formic acid in acetonitrile. The column used was a Waters Acquity UPLC BEH-C18 1.0 x 100.0 mm (Milford, MA). The gradient started at 1% B, then ramping to 99% B at T=10 min and holding for 3 minutes. There was a 4 min re-equilibration step. The flowrate was set to 150 µL/min and the sample injection volume was 8 µL. For back end mass spectrometry analysis, operating in positive ion mode, the source conditions were as follows: Gas temp = 200 ºC, Gas flow = 15 L/min, nebulizer pressure = 35 psi, Sheath gas temp = 325 ºC, Sheath gas flow= 9 L/min. Capillary voltage = 4000 V and nozzle voltage = 1500 V. Operating in negative ion mode, the source conditions were as follows: Gas temp = 200 ºC, Gas flow = 15 L/min, nebulizer pressure = 35 psi, Sheath gas temp = 325 ºC, Sheath gas flow= 9 L/min. Capillary voltage = 3000 V and nozzle voltage = 1500 V.

**Quantitative proteomics: sample preparation and analytical conditions.** Cell pellets were collected by centrifugation at 4°C at 6600 x g for 15 minutes. The pellet was washed once in 10 mL of 1X phosphate buffered saline (PBS; 137 mM NaCl, 2.7 mM KCl, 10 mM $Na_2HPO_4$, 1.8 mM $KH_2PO_4$ [pH 7.4]) followed by centrifugation at 5500 x *g* for 15 minutes at 4°C. The supernatant was decanted, and the cell pellets were frozen and stored at -80 °C until extraction and analysis. CPTF cell pellets were resuspended in Qiagen P2 Lysis Buffer (Qiagen, Hilden, Germany, Cat.#19052) to promote cell lysis. Proteins were precipitated with addition of 1 mM NaCl and 4 x vol acetone, followed by two additional wash with 80% acetone in water. The recovered protein pellet was homogenized by pipetting mixing with 100 mM Ammonium bicarbonate in 20% Methanol. Protein concentration was determined by the DC protein assay (BioRad, Hercules, CA). Protein reduction was accomplished using 5 mM tris 2-(carboxyethyl)phosphine (TCEP) for 30 min at room temperature, and alkylation was performed with 10 mM iodoacetamide (IAM; final concentration) for 30 min at room temperature in the dark. Overnight digestion with trypsin was accomplished with a 1:50 trypsin:total protein ratio. The resulting peptide samples were analyzed on an Agilent 1290 UHPLC system coupled to a Thermo Scientific Obitrap Exploris 480 mass spectrometer for discovery proteomics [2]. Briefly, peptide samples were loaded onto an Ascentis® ES-C18 Column (Sigma–Aldrich, St. Louis, MO) and separated with a 10 minutes LC gradient (10% Buffer A (98% H2O, 2% ACN) – 35% Buffer B (2% H2O, 98% ACN)). Eluting peptides were introduced to the mass spectrometer operating in positive-ion mode and were measured in data-independent acquisition (DIA) mode with a duty cycle of 3 survey scans from m/z 380 to m/z 985 and 45 MS2 scans with precursor isolation width of 13.5 m/z to cover the mass range. DIA raw data files were analyzed by an integrated software suite DIA-NN [3]. DIA-NN determines mass tolerances automatically based on first pass analysis of the samples with automated determination of optimal mass accuracies. The retention time extraction window was determined individually for all MS runs analyzed via the automated optimization procedure implemented in DIA-NN. Protein inference was enabled, and the quantification strategy was set to Robust LC = High Accuracy. Output main DIA-NN reports were filtered with a global FDR = 0.01 on both the precursor level and protein group level. Comparative proteomics analyses between two experimental conditions were performed by using an in-house python script. Differentially expressed proteins with statistical significance (Welch's t-Test with Benjamini-Hochberg FDR procedure [4]) were reported. Functional annotations for differentially expressed proteins were performed using eggNOG-mapper (v2) with default parameters [5]. Protein networks were visualized using EVenn [6].

**Iron uptake determination.** CPTF cell pellets were resuspended in 10 mL HEPEs buffer (25 mM, pH 7.5) and centrifuged at 5500 x *g* for 15 minutes at 4°C. Two additional wash steps were performed following the same procedure as above, discarding the supernatant after the final wash. Cell pellets were frozen at -20 °C until extraction. To prepare whole cell extracts, thawed cell pellets were resuspended in 2 mL of the HEPEs buffer with 0.1 mg/mL each of lysozyme, DNase, and EDTA. Resuspended pellets were incubated in the lysis buffer at room temperature for 15 minutes. Following incubation, cells were lysed by three rounds of sonication with 30 second breaks in between each round. Each round consisted of 5 second on, 5 second off cycles at 80 amp repeated 5 times. The sonication steps were carried out in a Coy anaerobic chamber (85% Ar, 5% $H_2$) The lysate was centrifuged at 5,500 x *g* for 30 minutes. A sample of the supernatant was saved for determination of Fe in the whole cell extract (cytoplasm + membrane). Protein concentrations of the whole cell extracts. determined using the Bradford assay [7].

Samples for metal analysis were diluted between 1:20 and 1:100 into 2 % (vol/vol) trace-grade nitric acid (VWR, Pennsylvania, USA). Samples were incubated at 37 °C temperature with shaking for at least 1 hr before centrifugation at 5,500 x *g* for 20 min to pellet debris. Samples were analyzed for Fe using an Agilent 7900 Inductively Coupled Plasma Mass Spectrometer (ICP-MS) fitted with MicroMist nebulizer, UHMI-spray chamber, Pt cones and an Octopole Reaction System (ORS) collision cell (Agilent Technologies, Santa Clara, CA). The external calibration standardd, IV-ICPMS-71A, was used to create a 10-point curve from 0 - 1000 pb for each element and the internal calibration standard, IV-ICPMS-71D, was added to each sample (Inorganic Ventures, Christiansburg, VA). The data was collected using 3-point peak pattern, 3 replicates, 100 sweeps/rep and various integration times (all >0.3 s) using either no gas or He gas in the ORS collision cell. Data was processed using the nearest internal standard by mass and fitted to a linear curve though the calibration blank.

**Enzyme activity assays.** Washed CPTF cell pellets were resuspended in 2 mL of the 50 mM potassium phosphate buffer (pH 7.0). The cell suspension (500 µL) was added to 4.5 mL of the assay buffer (50 mM potassium phosphate buffer [pH 7.0], 20 mM glucose, 1 mM nitrate). The assay mixture was incubated at 30°C while shaking at 150 rpm for 15 minutes. Following incubation, the OD600 of the assay mixture was determined. A subsample was taken and centrifuged at 18,000 x *g* for 5 minutes to separate the supernatant from the biomass. The supernatant was diluted 10-fold into sterile double deionized water. To determine [nitrate + nitrite], 50 µL of the diluted supernatant was added to 100 µL Greiss reagent [8] (4 g/100 mL) and 50 µL VCl3 (0.8 g/100 mL) and incubated at 30°C for 4 hours before measuring absorbance at 540 nm. To determine [nitrite], 100 µL of diluted supernatant was added to 100 µL of Greiss reagent and incubated at 30 °C for 15 minutes before measuring absorbance at 540 nm. The difference between [nitrate + nitrite] and [nitrite] was used to determine [nitrate]. Nitrate reductase activity was calculated with the following formula and normalized against OD600 values:

$$1\ unit\ of\ activity = \frac{\Delta([\text{nitrate}]_{\text{blank}} - [\text{nitrate}]_{\text{sample}})}{\text{time}}$$

One unit of activity is defined as one nmol of nitrate reduced per minute. Nitrite reductase assays were performed in the same manner as the nitrate reductase assays but with a different assay buffer composition: 50 mM potassium phosphate buffer (pH 7.0), 20 mM glucose, and 1 mM nitrite. Nitrite concentrations following 15 minutes of incubation were determined using the Greiss reagent as described above. Nitrite reductase activity was calculated with the following formula and normalized against OD600 values:

$$1\ unit\ of\ activity = \frac{\Delta([\text{nitrite}]_{\text{blank}} - [\text{nitrite}]_{\text{sample}})}{\text{time}}.$$

One unit of activity is defined as one nmol of nitrite reduced per minute.

**RNA extraction and cDNA preparation.** For RNA extraction, CPTF cultures were immediately chilled on ice for 10 min to stop growth. Cells were subsequently harvested by centrifugation (10 min, 7,500 rpm). Cells were resuspended in 0.3 mL lysis buffer (4 M guanidine thiocyanate, 0.83% N-lauryl sarcosine, pH5). After resuspension, 0.3 mL of acid-equilibrated phenol:chloroform 5:1 (pH 4.7) was added to each sample, and the samples were sonicated on ice (three 10 second pulses at amplitude 40 with 30 second rest periods in between). RNA was purified from the samples as previously described [9]. RNA was digested with TURBO DNAse (Ambion, Austin, TX) to remove contaminating genomic DNA. The quality of the RNA was evaluated by A260/A280. cDNA synthesis was performed using 1 µg purified RNA with the Affinity Script Quantitative reverse transcriptase PCR (qRT-PCR) cDNA synthesis kit (Agilent).

### SUPPLEMENTARY TABLES AND FIGURES

#### Table S1. MS transition states for metabolite analyses

| Compound | Polarity | Precursor ion (MS1) | Quantifier ion transition (MS2) | Qualifier ion transition (MS2) | Collision energy (V) |
|---|---|---|---|---|---|
| Tryptophan | Positive | 205.1 | 188 | 146.1 | 6/18 |
| Anthranilic acid | Positive | 138.06 | 65.1 | 92 | 42/26 |
| Chorismic Acid | Negative | 207 | 137.3 | 93.2 | 30/38 |
| 2,3-Dihydroxybenzoic Acid | Negative | 153 | 109.3 | 108.3 | 17/29 |

**Table S2. Quantitation range for analyzed metabolites**

| Compound | Lower limit (nM) | Upper limit (nM) |
|---|---|---|
| Tryptophan | 1000 | 50000 |
| Anthranilic acid | 200 | 10000 |
| Chorismic Acid | 10 | 1000 |
| 2,3-Dihydroxybenzoic Acid | 50 | 10000 |

### Table S3. qRT-PCR primer sequences

| Protein | GenBank ID | Gene name | Forward primer | Reverse primer |
|---|---|---|---|---|
| Iron-hydroxamate abc transporter substrate-binding protein | UIJ64690.1 | *feuA* | GTGTAGGTTTAGCGGCATGTA | TTGCTGGCACTGTATACTCTTT |
| Peptide mfs transporter | UIJ66957.1 | *mfs* | TCTTCTGGGCAGGATTTGAG | GGAACCAAGGTGTTGGAATTG |
| Isochorismatase family protein | UIJ68572.1 | *dhbZ* | CGTGAGCCAGTAGAGAGTATTG | CAGTTGGACGTTCTGCTAAATC |
| Isochorismate synthase dhbc | UIJ68570.1 | *dhbC* | CATTTGACCGCGAGTTCTTTAC | CAACTCCAGCTCCTGCATATAA |
| (2,3-dihydroxybenzoyl)adenylate synthase | UIJ68571.1 | *dhbY* | CAGTTCTTCCAGTAGCACACA | ACTACCACCAGTTGCCAATAC |
| Anthranilate synthase component i | UIJ67618.1 | *trpE* | TCCGGAAGTACACCTCTTACTC | CCGAATCATCTTCCCTGCATAC |
| Anthranilate phosphoribosyltransferase | UIJ67620.1 | *trpD* | AAATGTATAGAGCAGGGCTTCTT | GCAGTTTCACCTTTCGCTTTC |
| Tryptophan synthase subunit beta | UIJ67623.1 | *trpB* | GCGGTGGTGCAAAGATTTATT | TAGTGCCTGACCGATTGTATTG |
| Flavodoxin | UIJ64511.1 | *fld* | TGAATTACCATTTGAAGCGGAAG | CTGTATCACCCGATCCGAATAC |
| Nitrate reductase subunit beta | UIJ68393.1 | *narH* | ATGCATCGGTTGCCATACT | CAATGCCAGGTTTCGTTTCTAC |
| Sulfite reductase | UIJ67790.1 | *sir* | CGCCTGCAACGAAAGAAATAG | GCTAAATACGGTCGAGCTGATAA |
| Heme-degrading monooxygenase hmoa | UIJ69081.1 | *hmoA* | CGACGAAGAAGTTGTCGTAATG | CGCTCTATGTCCAGCTAAGT |
| Protein recA | UIJ64731.1 | *recA* | ATATCCACGTGGCCGTATTATC | CCACCTTGACGTTGTACTTCT |
| Nitrate reductase alpha subunit | UIJ68392.1 | *narG* | GCGTGGTAAAGGTGGATTTATTC | TTTCGATCTGGCCCGTATTT |
| Nitrate reductase gamma subunit | UIJ68395.1 | *narI* | CGTAATTGGCGGTCATGTTATG | CCGATGATAGAAGCAACACCT |

**Table S4. Geochemical parameters used to determine laboratory test values for experimentation with strain CPTF**

| Metal | [ORR Groundwater Median] (µM)[1] | [COMM] (µM) |
|---|---|---|
| **Al** | 3385 (215-20700) | 500[2] |
| **U** | 110 (40-576) | 50[2] |
| **Mn** | 1202 (446-3150) | 50[2] |
| **Ni** | 59 (19-157) | 50 |
| **Co** | 8 (2-30) | 15 |
| **Cu** | 1 (0.2-15) | 5 |
| **Fe** | 1 (0.6-9.9) | 5 |
| **Cd** | 1 (0.6-10) | 2.5 |

*[1]Median values for six Area 3 wells, described previously by Thorgersen et al. (2019).The range is given in parentheses. [2]Concentrations used were limited by the solubility of the metal at neutral pH.*

**Table S5. Error bars (± SD) for Figure 1A**

| Time (h) | 0 | 3 | 4 | 5 | 6 | 7 | 8 | 9 | 10 | 13 | 14 |
|---|---|---|---|---|---|---|---|---|---|---|---|
| **Control** | 0.000 | 0.001 | 0.006 | 0.008 | 0.004 | 0.021 | 0.014 | 0.020 | 0.019 | 0.003 | 0.006 |
| **COMM** | 0.000 | 0.004 | 0.004 | 0.004 | 0.004 | 0.012 | 0.016 | 0.005 | 0.018 | 0.006 | 0.007 |
| **Al** | 0.000 | 0.011 | 0.017 | 0.026 | 0.049 | 0.064 | 0.067 | 0.075 | 0.067 | 0.021 | 0.016 |
| **U** | 0.000 | 0.005 | 0.009 | 0.007 | 0.009 | 0.015 | 0.029 | 0.019 | 0.008 | 0.008 | 0.023 |
| **Mn** | 0.000 | 0.001 | 0.002 | 0.003 | 0.013 | 0.020 | 0.019 | 0.038 | 0.017 | 0.011 | 0.003 |
| **Ni** | 0.000 | 0.002 | 0.003 | 0.017 | 0.007 | 0.010 | 0.017 | 0.012 | 0.005 | 0.003 | 0.009 |
| **Co** | 0.000 | 0.002 | 0.003 | 0.008 | 0.010 | 0.010 | 0.006 | 0.023 | 0.034 | 0.016 | 0.012 |
| **Cu** | 0.000 | 0.014 | 0.019 | 0.020 | 0.029 | 0.026 | 0.017 | 0.010 | 0.009 | 0.018 | 0.023 |
| **Fe** | 0.000 | 0.010 | 0.011 | 0.021 | 0.034 | 0.053 | 0.052 | 0.041 | 0.036 | 0.013 | 0.010 |
| **Cd** | 0.000 | 0.010 | 0.021 | 0.019 | 0.019 | 0.024 | 0.021 | 0.030 | 0.020 | 0.011 | 0.014 |

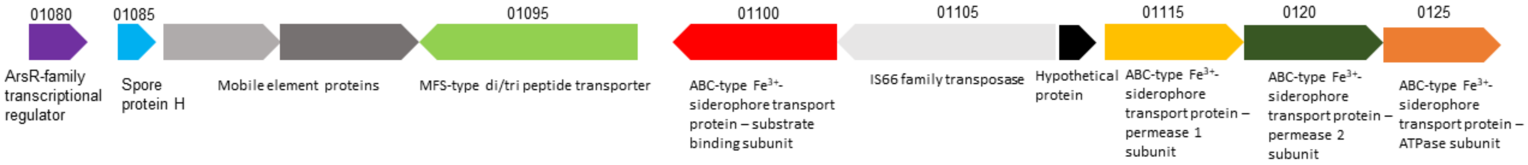


**Figure S1. Genomic location of the predicted bacillibactin exporter.** The MFS-type di/tri-peptide transporter (RefSeq: LW858_01095) is predicted to export bacillibactin during COMM exposure. The genomic context of this gene is shown. Genes are labeled with RefSeq locus tags (when available) and annotations.

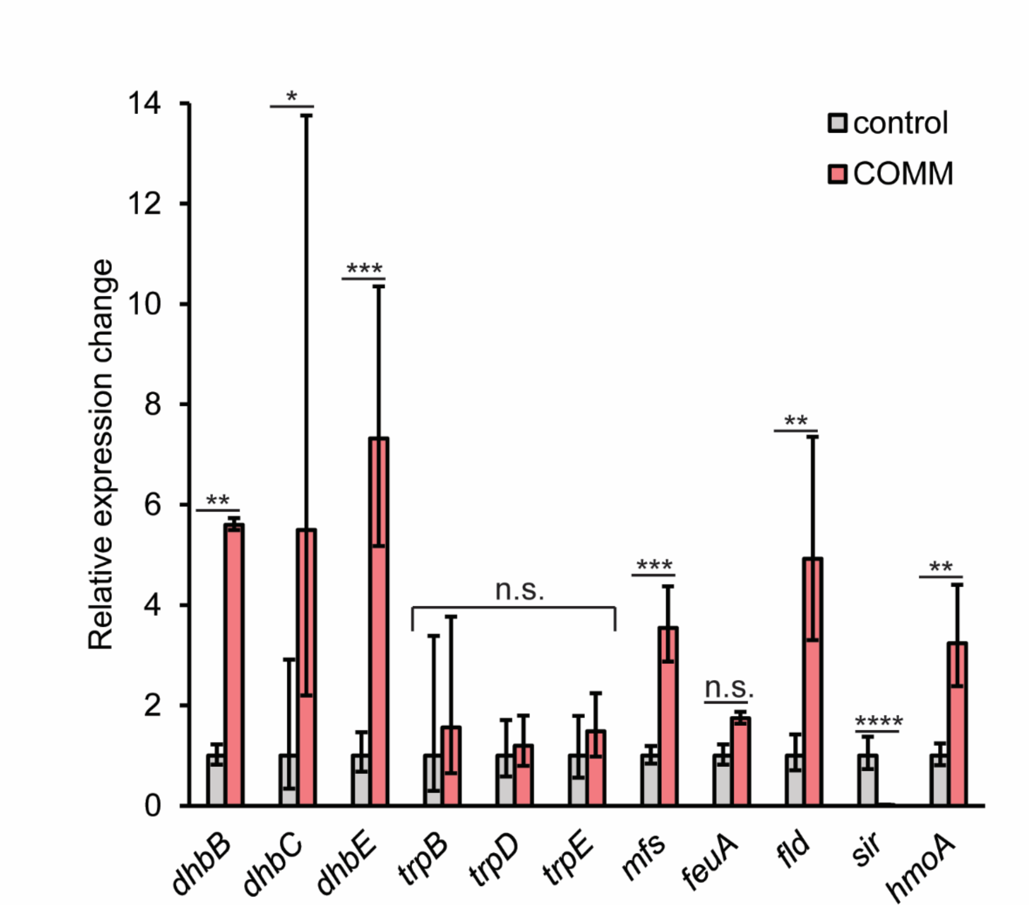


**Figure S2. qRT-PCR validation of protein expression changes.** Relative changes in expression of select genes during (red bars) and without (grey bars) COMM exposure. Experiments were performed in triplicate and error bars represent ±SD. * $p < 0.05$, ** $p < 0.01$, *** $p < 0.001$, **** $p < 0.0001$, n.s. difference between expression levels is non-significant ($p > 0.05$).

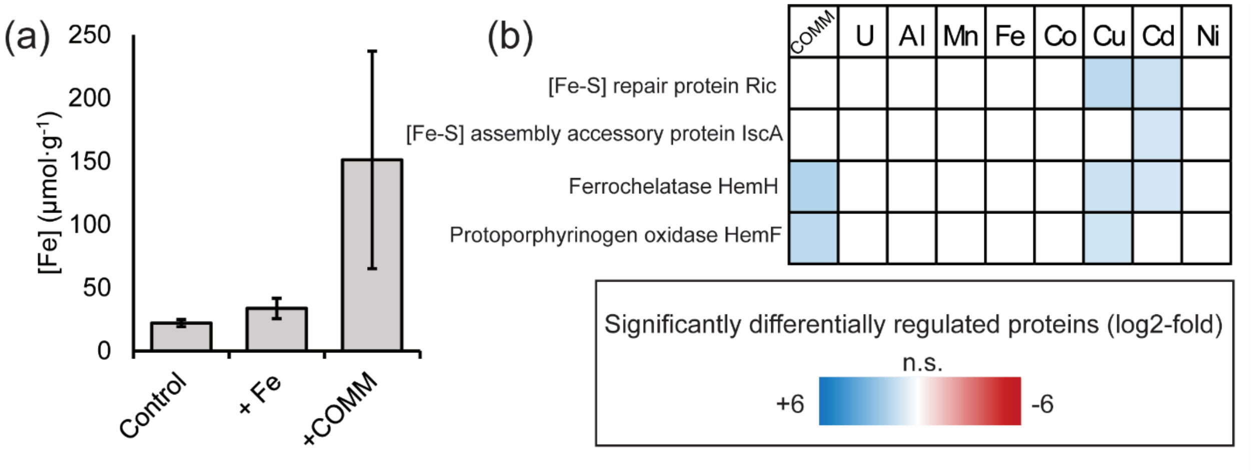


**Figure S3. Experimental insights into possible mechanism for iron starvation response. (a)** Total iron concentrations measured in whole cell extracts of unamended cultures (control), cultures amended with 5 µM $Fe^{2+}$ (+ Fe), and cultures amended with COMM (+COMM). Values are normalized against protein measurements. Bars are the average of three replicates and error bars represent ±SD. **(b)** Expression patterns of proteins involved in iron cofactor biosynthesis. Heat map displays average (n= 3 replicates) $log_2$-fold expression changes of individual proteins across the different treatment conditions relative to the control (left-to-right: COMM, U, Al, Mn, Fe, Co, Cu, Cd, Ni). The heat map scale (displayed at $log_2$-fold expression changes relative to the control) is at the bottom of the image.
